# Anti-oxidant Response of Lipidom Modulates Lipid Metabolism in *Caenorhabditis elegans* and in OxLDL-Induced Human Macrophages by Tuning Inflammatory Mediators

**DOI:** 10.1101/2022.11.14.516538

**Authors:** Acharya Balkrishna, Vivek Gohel, Nishit Pathak, Rani Singh, Meenu Tomer, Malini Rawat, Rishabh Dev, Anurag Varshney

## Abstract

Atherosclerosis is the main pathological process of most cardiovascular diseases. It can begin early in life and may remain latent and asymptomatic for an extended period before its clinical manifestation. Lipidom, an ayurvedic prescription medicine, contains five herbal constituents with reported anti-inflammatory, anti-oxidant and lipid lowering properties. The present study is aimed to characterize the pharmacological potentials of Lipidom. The phytochemical analysis of Lipidom was performed on high performance liquid chromatography (HPLC) platform. Lipidom was evaluated for cytosafety, NF-κB activity, IL-1β and MCP-1 levels, modulation of NLRP3 pathway, ROS generation, lipid accumulation and gene expression in oxidized LDL stimulated THP1 macrophages. Furthermore, assessment of Lipidom was also done in the in-vivo *Caenorhabditis elegans* model. Analysis of brood size, % adult, lipid accumulation, triglyceride levels, MDA formation, SOD-3 levels and gene expression was performed in *C. elegans*. Lipidom treatment significantly reduced the inflammatory markers, lipid accumulation, oxidative stress and normalized genes involved in atherosclerosis development in THP1 macrophages. Lipidom treated *C. elegans* showed a significant decline in the lipid accumulation and oxidative stress. Lipidom showed a multifaceted approach in modulating the mediators responsible for development and progression of atherosclerosis.

## Introduction

Atherosclerosis is a chronic inflammatory disease which occurs in the arterial wall. It is generally characterized by the formation of plaques containing lipids, immune cells and connective tissue in the intima of arteries. Inside the arteries the immune cells are activated due to persistent inflammatory conditions or a failure in the resolution of inflammation. This leads to the development of chronic inflammation which is a hallmark of cardiovascular diseases (CVD) [1]. It is estimated that by the year 2030 nearly 23.6 million individuals will die from CVD annually [2]. One of the major risk factors for the development of atherosclerosis is occurrence of lipid metabolism disorders like dyslipidaemia and hypercholesterolemia [3]. The monocytes play an important role in the progression of the disease. Modified lipoproteins like oxidized low density lipoproteins (OxLDL) recruit the monocyte to the vascular intima where they get differentiated to macrophages to engulf the OxLDL and turn to foam cells. The accumulation of these foam cells in the subendothelial space primarily leads to the initiation and progression of atherosclerosis [4].

The current pharmacotherapy to prevent the development of atherosclerosis aims to lower the lipid levels. Statins forms the most widely prescribed drug class for management of lipid levels but the chances of sub-optimal dosing and drug discontinuation occurs in nearly 20% of the patients due to the development of side effects like elevation of blood glucose and new-onset diabetes [3]. Herbal drugs like Lipidom are being increasingly used for their anti-oxidant, lipid lowering and anti-inflammatory properties. Lipidom is composed from the extracts of five herbs namely Arjuna (*Terminalia arjuna*), Lehsun (*Allium sativum*), Dalchini (*Cinnamomum verum*), Guggul (*Commiphora wightii*), Lauki (*Lagenaria siceraria*). The description of the medicinal properties of these plants have been described in the classical ayurvedic text Bhavprakash Nighantu (BPN) (Table 1).

**Table 1.** Herbal composition of Lipidom.

| Sl. No. | Common Name | Latin Name | Part used | Book Ref. | Page no. | Qty. (mg/tablet) |
| --- | --- | --- | --- | --- | --- | --- |
| 1. | Arjuna Dry Extract | *Terminalia arjuna* | Bark | B.P.N | 523-524 | 170 |
| 2. | Lehsun Dry Extract | *Allium sativum* | Bulb | B.P.N | 131-134 | 40 |
| 3. | Dalchini Dry Extract | *Cinnamomum verum* | Stem bark | B.P.N | 226 | 40 |
| 4. | Guggul Dry Extract | *Commiphora wightii* | Exuate gum resin | B.P.N | 205 | 40 |
| 5. | Lauki Dry Extract | *Lagenaria siceraria* | Whole Plant | B.P.N | 681-682 | 170 |

The present study investigated the anti-atherogenic properties of Lipidom. A thorough phytochemical evaluation of Lipidom was carried out via HPLC followed by assessment of its pharmacological effects. The in-vitro analysis of Lipidom was done by evaluation of parameters like cytosafety, NF-κB activity, IL-1β and MCP-1 levels, modulation of NLRP3 pathway, ROS generation, lipid accumulation and gene expression. Furthermore, Lipidom evaluation was also performed in the in-vivo *Caenorhabditis elegans* model. Assessment of brood size, % adult, lipid accumulation, triglyceride levels, MDA formation, SOD-3 levels and gene expression was performed in *C. elegans* treated with Lipidom. Atorvastatin [5] and a combination of leucine-nicotinic acid [3] was used as the positive control for the in-vitro and in-vivo analysis respectively.

## Materials and Methods

### Reagents

Lipidom (batch # 1LPD-210044) was sourced from Divya Pharmacy, India. The standards for HPLC analysis namely Protocatechuic acid, Corilagin, Guggulsterone E and Guggulsterone Z were obtained from Natural remedies, India; Gallic acid and Ellagic acid from Sigma-Aldrich, USA, and Cinnamic acid from SPL, USA. Reagents namely antibiotic-antimycotic solution, Malondialdehyde (MDA), Trichloroacetic acid (TCA), LPS, Paraquat, and 2’,7’-Dichlorofluorescin diacetate (H2DCFDA) were procured from Sigma-Aldrich, USA. Heat-inactivated FBS and Alamar blue were obtained from HiMedia, India. RPMI 1640 was bought from Gibco, USA. Chemicals, 2-thiobarbituric acid (TBA), Atorvastatin, leucine, and nicotinic acid were obtained from TCI chemicals, India. Phorbol 12-myristate 13-acetate (PMA) was purchased from Alfa Aesar, UK. LipidSpot 610 reagent was procured from Biotium, USA. OxLDL, TNF-α, Verso cDNA synthesis kit, TriZol, CellROX Green reagent, Pierce BCA protein assay kit was obtained from Thermo Fisher Scientific, USA. The RNeasy mini kit was purchased from Qiagen, Germany. The LDH and triglyceride estimation reagents were obtained from Erba Mannheim, Germany PowerUp SYBR Green Master Mix was procured from Applied Biosystems, USA. Human IL-1β ELISA kit was obtained from BD Biosciences. CB-Protein Assay kit and MCP-1 ELISA kit from G-Biosciences, India. QUANTI-Blue reagent was purchased from InvivoGen, USA.

### Phytometabolite analysis of Lipidom

Lipidom powder (500 mg) was diluted with 10 ml methanol: water (80:20) and sonicated for 30 min. This solution was centrifuged at 10000×g for 5 min and filtered by 0.45 μm nylon filter. This solution was further used for the phytochemical analysis. Analysis was performed by Prominence-XR UHPLC system (Shimadzu, Japan) equipped with Quaternary pump (NexeraXR LC-20AD XR), DAD detector (SPD-M20 A), Auto-sampler (Nexera XR SIL-20 AC XR), Degassing unit (DGU-20A 5R) and Column oven (CTO-10 AS VP). The elution was carried out at a flow rate of 1.0 ml/min using gradient elution of mobile phase A (0.1% acetic acid in water and mobile Phase B (Acetonitrile). The experiment was performed on Shodex C18-4E (4.6 x 250 mm, 5 μm) column. Gradient programming of the solvent system for mobile phase B was set as 5 % for 0-10 min, 5-10 % from 10-20 min, 10-15 % from 20-30 min, 15 % from 30-40 min, 15-40 % from 40-55 min, 40-60 % from 55-60 min, 60-90 % from 60-65 min, 90-5% from 65-68 min and 5 % from 68-70 min with a flow rate of 1.0 mL/min. 10 μl of standard and test solution were injected and column temperature was maintained at 35 °C. The analysis of gallic acid, protocatechuic acid, corilagin, ellagic acid, and cinnamic acid was performed at a wavelength of 270 nm and of guggulsterone E and guggulsterone Z at 240 nm.

### Cell culture maintenance

The human monocytic cells THP-1 were procured from the ATCC licensed cell repository, National Centre for Cell Science, India. The reporter cell line THP1-Blue NF-κB was obtained from InvivoGen, USA. THP-1 cells were cultured in RPMI 1640 supplemented with 10 % FBS and 1 % antibiotic-antimycotic solution. The THP1-Blue NF-κB cells were cultured as per the manufacturer’s instructions. The cultured cells were maintained at 37 °C and 5% CO_2_ in a humidified incubator and used within 5 passages after revival.

### Cytosafety analysis of Lipidom and OxLDL treated THP1 cells

The THP1 monocytes were seeded at a density of 1×10^5^ cells/well in a 96-well plate in presence of 60 ng/ml PMA for differentiation to macrophages. The cells were incubated for 72 hr at 37 °C and 5% CO_2_ after which the cells were washed twice with DPBS and incubated with Lipidom (1-1000 μg/ml), Atorvastatin (40, 80, and 160 μM), and DMSO (1% v/v) as vehicle control. The cells were further incubated for 96 hr post which the cells were washed twice with DPBS and incubated with 100 μl of Alamar blue cell viability reagent. The plates were evaluated for fluorescence at Ex. 560/ Em. 590 nm on infinite 200Pro (Tecan, Switzerland) plate reader. Data were presented as mean ± SEM (n = 3). The cytosafety analysis of OxLDL (25, 50, and 100 μg/ml) induced differentiated THP1 macrophages was performed via assessment of the released LDH (U/L) levels post 72 hr incubation of cells with OxLDL. After incubation the media supernatant was collected and the levels of LDH were evaluated by Erba 200 biochemical analyzer (Erba Mannheim, Germany). Untreated cells (UC) were used as normal control.

### Assessment of TNF-*α* induced NF-κB activity

THP1-Blue NF-κB reporter cells were used for monitoring the level of NF-κB activation. Briefly, cells with density of 5×10^5^/ml were seeded in 96-well plate and co-treated with 10 ng/ml TNF-α and Lipidom (10, 30, and 100 μg/ml) or Atorvastatin (40 μM) for 24 hr post which the evaluation of Secreted Embryonic Alkaline Phosphatase (SEAP) was done by QUANTI-Blue Solution as per the manufacturer’s instructions. The optical density was read at 630 nm using an Envision (PerkinElmer, USA) multimode plate reader. Data were presented as mean ± SEM (n = 3).

### Assessment of OxLDL-induced IL-1β and MCP-1 release in THP1 macrophages

THP1 monocytes were plated at a density of 1×10^5^ cells per well. The cells were differentiated to macrophages and pre-treated with Lipidom (10, 30, and 100 μg/ml) or Atorvastatin (40 μM) for 24 hr. Post incubation, the cells were washed and stimulated with 25 μg/ml OxLDL in presence of Lipidom (10, 30, and 100 μg/ml) and Atorvastatin (40 μM) for 72 hr. The cell supernatant was then collected and assessed for IL-1β and MCP-1 release by sandwich ELISA as per the manufacturer’s instructions. Data were presented as mean±SEM (n=3).

### NLRP3 assay

PMA (20 ng/ml) differentiated THP1 macrophages (1×10^5^/ well) were grown in 96-well culture plate in the RPMI media containing 10 % FBS. Cells pre-treated with Lipidom (10, 30, and 100 μg/ml) or Atorvastatin (40 μM) for 24 hr. Thereafter, media was removed and cells were further primed with 100 ng/ml LPS for 4 hr in complete media. After completion of incubation time, cells were washed with PBS and further co-treated with various concentrations of Lipidom along with ATP (5 mM) in the plain RPMI media for 45 min. The supernatants were collected and evaluated for IL-1β cytokine release using ELISA as per the manufacturer’s protocol. Data were presented as mean±SEM (n=3).

### Microscopy based assessment of lipid accumulation and ROS generation

THP1 monocytes were plated at a density of 5×10^4^ cells/well on 8-well cell culture slide. The cells were differentiated to macrophages and pre-treated with Lipidom (100 μg/ml) for 24 hr. Post incubation, the cells were washed and stimulated with 25 μg/ml OxLDL in presence of Lipidom (100 μg/ml) for 72 hr. Thereafter, cells were washed and simultaneously incubated with 5μM of CellROX green reagent and LipidSpot 610 (1:1000) lipid droplet stain for 15 min. The cells were washed and mounted with ProLong Diamond antifade mountant with DAPI (Invitrogen, USA). Microscopy was performed by Olympus BX43 microscope equipped with Mantra imaging platform (Perkin Elmer, USA) and further processed on Inform 2.2 software suite (Perkin Elmer, USA).

### Evaluation of ROS levels in OxLDL-induced THP1 macrophages

ROS levels were detected by the H2DCFDA probe dye. Briefly, after treatment cells were lysed, centrifuged and the supernatant was incubated with H2DCFDA (20 μM) at 37 °C for 1 hr. The plates were read at Ex. 495/Em.523 nm by Envision multimode plate reader and the fluorescence values were normalized with protein concentration. Data were presented as mean±SEM (n=3).

### Evaluation of MDA levels in OxLDL-induced THP1 macrophages

The MDA content from the lysate of treated THP1 cells was measured by the TBA-TCA method. The optical density was read at 532 nm by through infinite 200Pro (Tecan, Switzerland) plate reader and the amount of MDA present in the samples was determined from the standard curve. The data was further normalized with protein concentration. Data were presented as mean±SEM (n=3).

### Gene expression assessment of OxLDL-induced THP1 macrophages

Evaluation of the mRNA expression levels of various genes was done by qRT-PCR. The total RNA was extracted by RNeasy mini kit following the manufacturer’s protocol. The cDNA synthesis was done using the Verso cDNA synthesis kit. cDNA samples were mixed with the PowerUp SYBR Green Master Mix and RT-PCR was performed using qTOWER3 G (Analytik-Jena, Germany). The qRT-PCR cycling parameters included an initial denaturation of 95 °C for 10 min and a primer extension at 95 °C for 15 sec and 60 °C for one minute with 40 cycles. Ct values were obtained, and relative expression 2^(-ΔΔCt)^ was calculated and analyzed for changes in mRNA expression. Primers used for the study are mentioned in Table 2. RPL-13 gene was used as the housekeeping gene. Data were presented as mean±SEM (n=3).

**Table 2.**
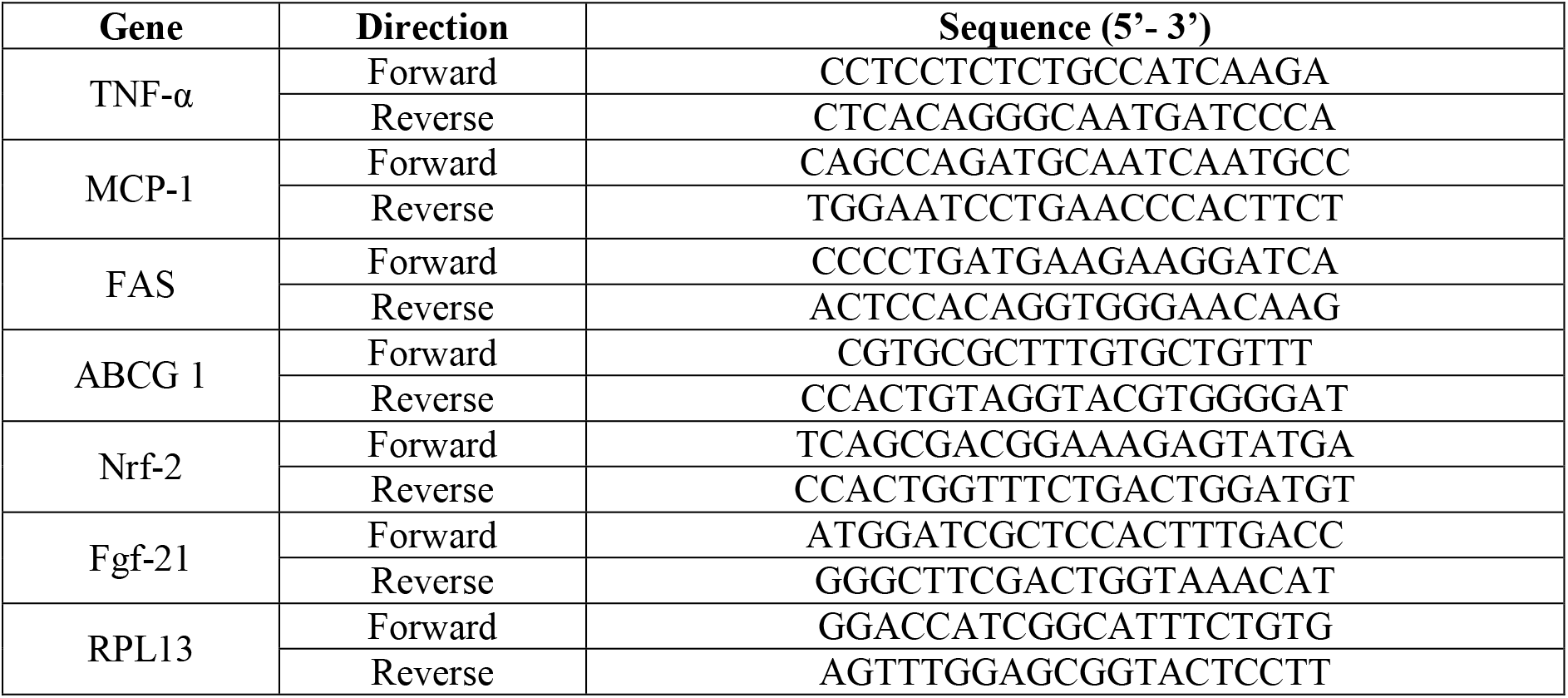
Primers used for mRNA expression analysis of THP1 cells.

### Maintenance of *Caenorhabditis elegans*

The strains N2 (wild type), CF1553 (muls84[pAD76(sod-3::GFP)]), and *Escherichia coli* OP50 were obtained from the Caenorhabditis Genetics Center at the University of Minnesota, USA. All strains were maintained in nematode growth medium (NGM) agar plates at 20 °C. The worms were seeded with living *E. coli* OP50 as a food source. A synchronization technique was used to separate the eggs from the worms by treatment with an alkaline hypochlorite solution. The hatched eggs released the L1 larvae which have been used for various treatments. The L1 nematodes were exposed to different concentrations of Lipidom (3, 10, and 30 μg/ml) or Nicotinic acid (10nM)-Leucine (0.5mM) (NA-Leu), transferred to an NGM plate seeded with *E. coli* OP50. A treatment scheme and parameters analyzed on *C. elegans* are mentioned in Figure 1.

**Figure 1.**
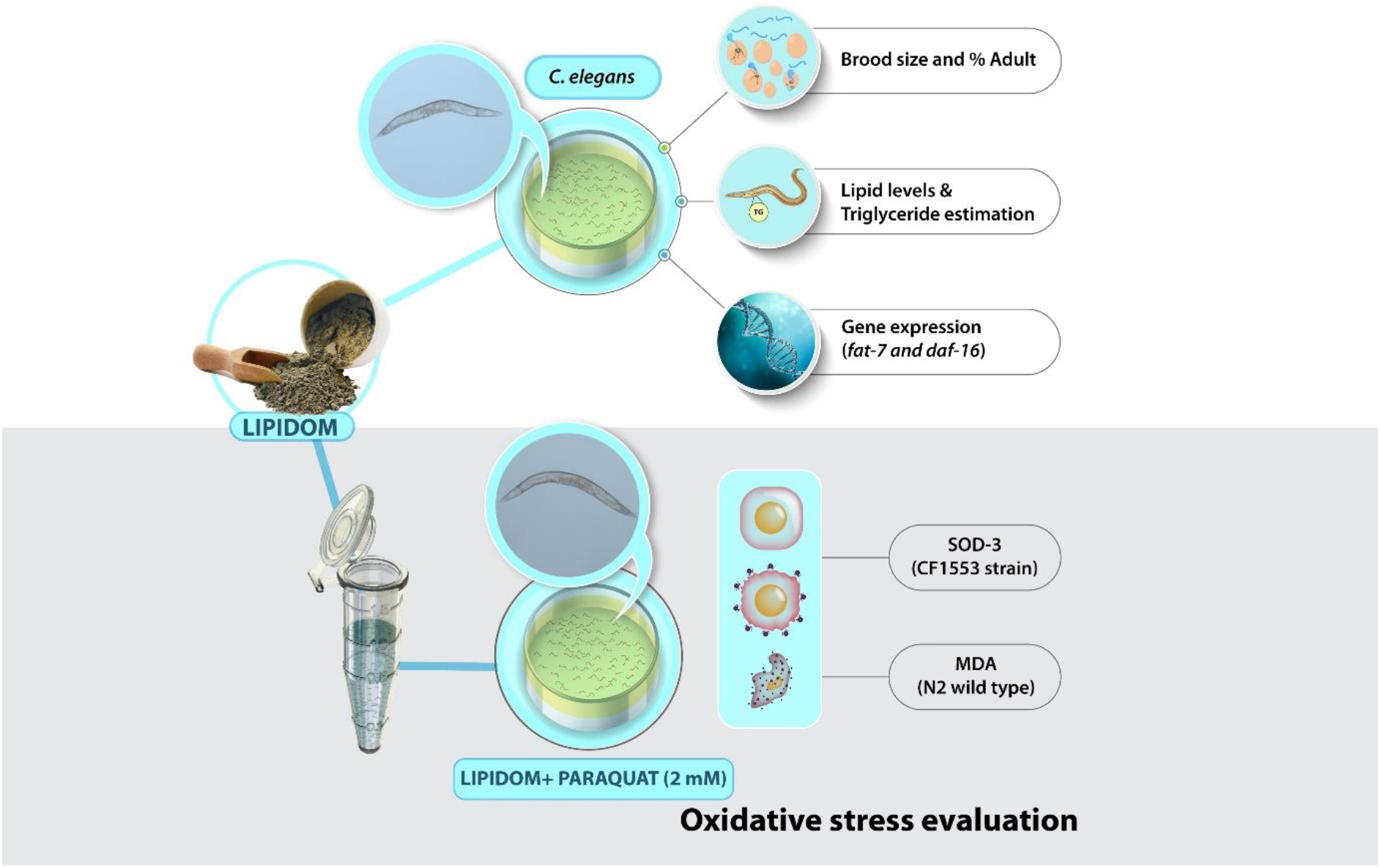
Schematic of *C. elegans* treatment and parameters evaluated.

### Assessment of Brood size and % Adult of *C. elegans*

The nematodes were treated as described above and were maintained on NGM/E. coli OP50 plates until they reached larval stage L4. To evaluate progeny size, one nematode from each treated group was transferred to a NGM plate with *E. coli* OP50 supplemented with Lipidom (3,10, and 30 μg/ml) or NA –Leu. The total number of progenies was counted. The determination of % Adult was done by counting the number of L1 stage worms on Day 1 and the number of adults developed at Day 7. Counting was done using the ZEISS Stemi 305 stereo microscope (Carl Zeiss, Germany). Data were presented as mean±SEM (n=5). Data were presented as mean±SEM (n=5).

### Assessment of triglyceride accumulation in *C. elegans*

The nematodes were treated as described above and were maintained on NGM/E. coli OP50 plates till Day 7. On Day 7 worms were washed off from the plates and centrifuged at 600×g for 1 min. Worms were washed rigorously and placed in M9 buffer and passed through three cycles of freeze-thaw. The lysates were sonicated for 1 min with the Ultra probe sonicator and then centrifuged at 14000×g for 15 min. The supernatant was collected and analyzed for triglyceride levels by Erba 200 biochemical analyzer (Erba Mannheim, Germany). Protein concentration was determined using the CB-Protein Assay kit. The obtained values of triglyceride levels were further normalized with protein concentration. Data were presented as mean±SEM (n=3).

### Assessment of lipid levels in *C. elegans*

The nematodes were treated as described above and were maintained on NGM/E. coli OP50 plates till Day 7. On day 7, ten adult worms from each group were picked and transferred into 96-well plate. Worms were washed rigorously with M9 buffer to remove the *E.coli*. After washing, LipidSpot 610 lipid droplet stain (1:1000) prepared in M9 buffer was added and the plate was kept in dark for 30 min on shaking. After 30 minutes, the plate was washed rigorously with M9 buffer. After washing worms were lyzed with 200 μl DMSO. The supernatant was collected on 96-well black plate for lipid levels at Ex.592nm/Em. 638 nm through infinite 200Pro (Tecan, Switzerland) plate reader. Protein concentration was determined using the CB-Protein Assay kit. The obtained values of lipid levels were further normalized with protein concentration. Data were presented as mean±SEM (n=3).

### Gene expression assessment of Lipidom treated *C. elegans*

The nematodes were treated as described above and were maintained on NGM/*E. coli* OP50 plates till Day 7. On Day 7, worms were washed off from the plates and centrifuged at 600×g for 1 min. Worms were washed rigorously and placed in TRIzol. After 5 freeze-thaw cycles RNA was isolated as per the manufacturer’s protocol. Subsequent cDNA preparation and qRT-PCR was performed as mentioned before. Primers used for the study are mentioned in Table 3. Act-1 was used as the housekeeping gene.

**Table 3.**
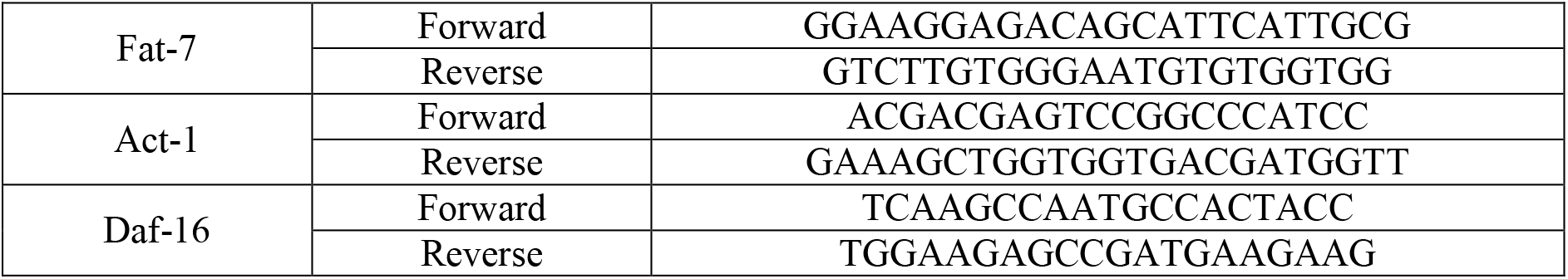
Primers used for mRNA expression analysis in *C. elegans*.

### Assessment of MDA levels in *C. elegans*

The L1 nematodes were exposed to different concentrations of Lipidom (3, 10, and 30 μg/ml) or NA-leu (10nM-0.5mM) for 24 hr. After incubation, the nematodes were transferred to an NGM plate seeded with *E. coli* OP50 with or without Paraquat (2 mM) supplementation along with Lipidom (3, 10, and 30 μg/ml) or NA-leu (10nM-0.5mM). On Day 7 worms were washed off from the plates and centrifuged at 600×g for 1 min. Worms were washed rigorously and placed in M9 buffer and passed through three cycles of freeze-thaw. The lysates were sonicated for 1 min with the Ultra probe sonicator and then centrifuged at 14000×g for 15 min. The MDA content from the lysate of treated nematodes was measured by the TBA-TCA method. The optical density was read at 532 nm through infinite 200Pro (Tecan, Switzerland) plate reader and the amount of MDA present in the samples was determined from the standard curve. The data was further normalized with protein concentration. Data were presented as mean±SEM (n=3).

### Assessment of SOD-3 activity in *C. elegans* (CF1553 strain)

The CF1553 (muls84[pAD76(sod-3::GFP)]) strain was maintained in nematode growth medium (NGM) agar plates at 20 °C. A synchronization technique was used to separate the eggs from the worms by treatment with an alkaline hypochlorite solution. The hatched eggs released the L1 larvae, used for exposure to various treatments. The L1 nematodes were exposed to different concentrations of Lipidom (3, 10, and 30 μg/ml) or NA-leu (10nM-0.5mM) for 24 hr. After incubation, the nematodes were transferred to an NGM plate seeded with *E. coli* OP50 with or without Paraquat (2 mM) supplementation along with Lipidom (3, 10, and 30 μg/ml) or NA-leu (10nM-0.5mM). On Day 7, the worms were flushed from plate, washed twice with M9 buffer and resuspended in M9 buffer. Afterwards, 100 μl of worm suspension was transferred to a 96-well black plate. The plate was read at Ex.480 nm/ Em.510 nm through infinite 200Pro (Tecan, Switzerland) plate reader. The worms were lysed and protein concentration was determined using the CB-Protein Assay kit. The obtained values were further normalized with protein concentration. Data were presented as mean±SEM (n=3).

### Data analysis

Statistical analysis was performed using one-way ANOVA with Dunnett’s multiple comparisons post-hoc test. Data were analyzed using GraphPad Prism 7 (GraphPad Software, USA). Results were considered to be statistically significant at a probability level of p < 0.05.

## Results

### Phytometabolite analysis of Lipidom

The quantitative analysis of phytochemicals using standard marker compounds (Figure 2A, 2B) confirmed the presence of Gallic acid (RT: 6.90 min), Protocatechuic acid (RT: 14.54 min), Corilagin (RT: 32.08 min), Ellagic acid (RT: 45.52 min), Cinnamic acid (RT: 56.40 min), Guggulsterone E (RT: 66.49 min), Guggulsterone Z (RT: 67.55 min) in Lipidom. The quantification of phytochemicals using reference standards is mentioned in Table 3.

**Figure 2.**
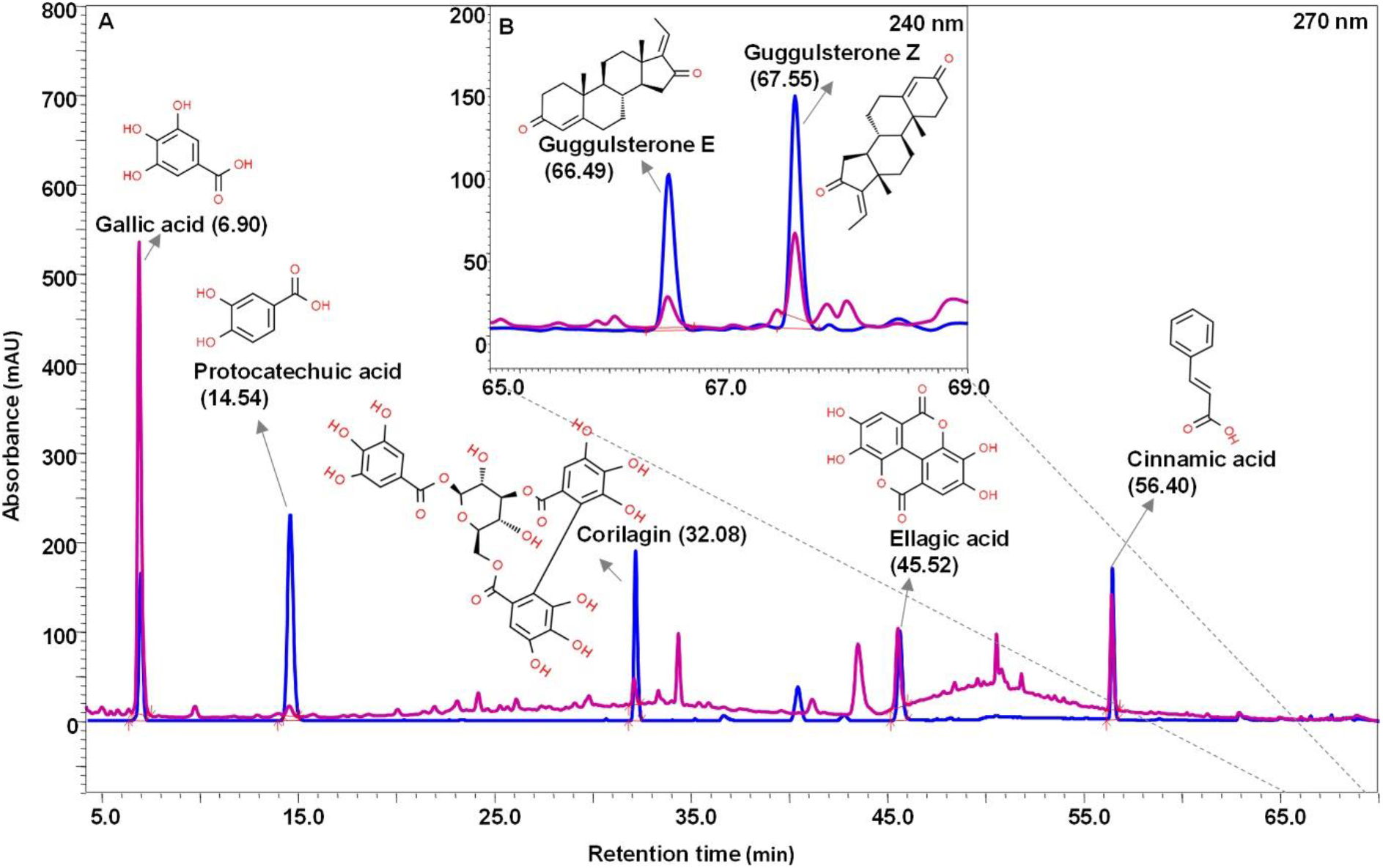
Phytometabolite analysis of Lipidom by HPLC. **A)** Overlap chromatogram of standard mix (blue line) and Lipidom (pink line) from 5 min to 70 min at 270 nm. **B)** Overlap chromatogram of standard mix and Lipidom tablet from 65 min to 69 min at 240 nm.

**Table 3.** Phytochemical constituents present in Lipidom. HPLC based quantification of phytochemicals in Lipidom

| Sr.No. | Compound | Quantity ($\mu\text{g}/\text{mg}$) |
| --- | --- | --- |
| 1 | Gallic acid | 4.537 |
| 2 | Protocatechuic acid | 0.066 |
| 3 | Corilagin | 0.189 |
| 4 | Ellagic acid | 1.280 |
| 5 | Cinnamic acid | 0.394 |
| 6 | Guggulsterone E | 0.079 |
| 7 | Guggulsterone Z | 0.151 |

### Cytosafety analysis

Lipidom was observed to be cytosafe upto 1000 μg/ml concentration on THP1 macrophages. Atorvastatin treatment at 80 and 160 μM concentration showed a significant (p <0.001) decline in cell viability (Figure 3A). The physiologically relevant Lipidom (10, 30, and 100 μg/ml) concentrations were used for further analysis. Similarly, Atorvastatin (40 μM) was used for other evaluations. The dose of OxLDL (25 μg/ml) was selected based on the released LDH (U/L) levels of OxLDL induced cells (Figure 3B).

**Figure 3.**
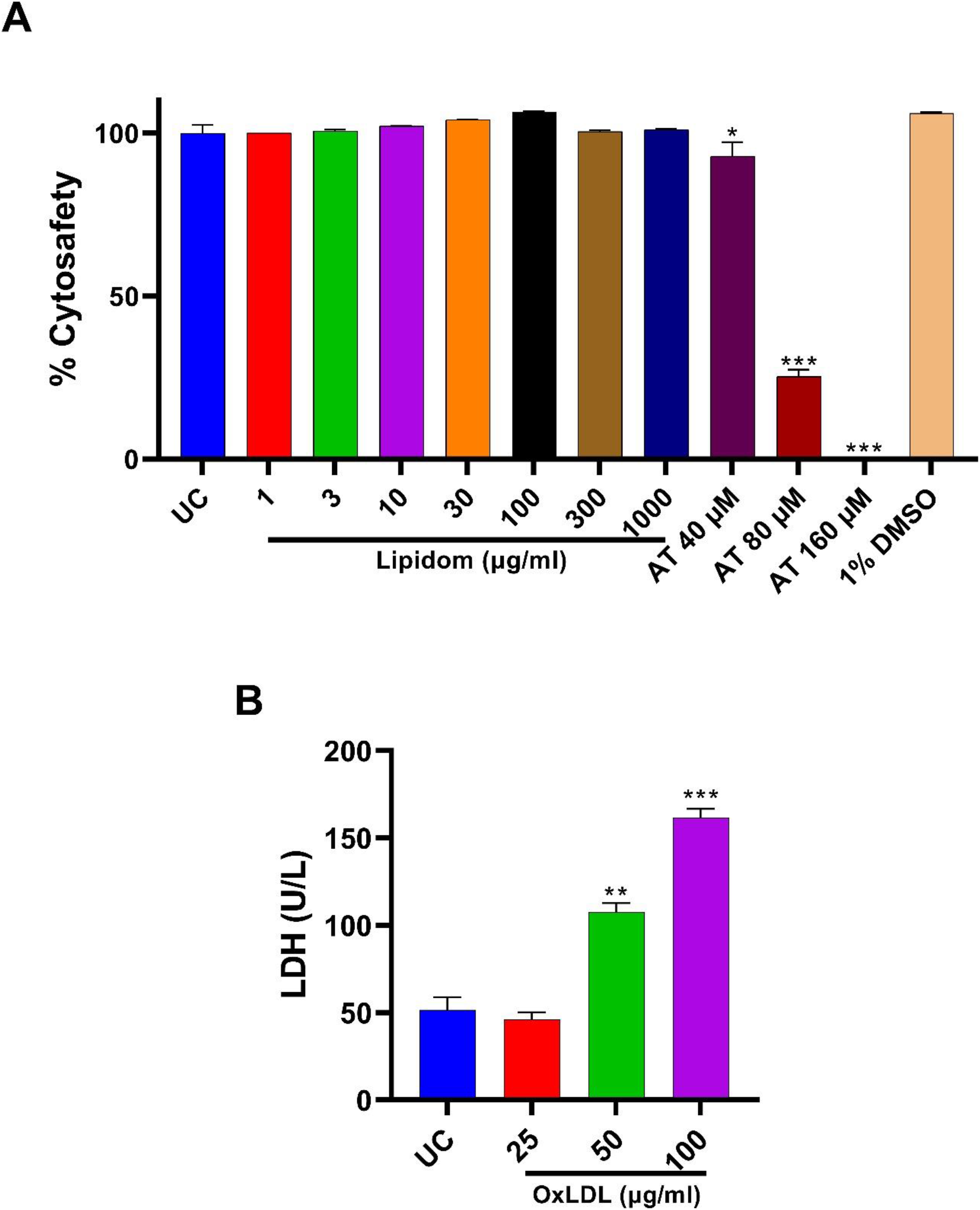
Cytosafety analysis. **A)** Cytosafety of Untreated cells (UC), Lipidom (1-1000 μg/ml) was performed on THP1 macrophages for 96 hr along with Atorvastatin (40-160 μM) and Vehicle control (1% DMSO v/v). **B)** LDH (U/L) levels of OxLDL (25, 50, and 100 μg/ml) on THP1 macrophages.

### Lipidom decreased inflammation by reduction of NFκB activity, IL-1β, MCP-1 release and modulation of NLRP3 inflammasome pathway

TNF-α, is a major proinflammatory cytokine which activates NFκB which aids in the release of inflammatory cytokines. Inflammatory mediators namely IL-1β, MCP-1 furthers the atherosclerotic plaque formation by migration, infiltration, proliferation, and differentiation of immune cells like monocytes and macrophages which leads to the development of chronic inflammation [6]. Lipidom (10, 30, and 100 μg/ml) treatment significantly (p <0.001) decreased the TNF-α induced NFκB activity and also inhibited the IL-1β, MCP-1 release from the OxLDL (25 μg/ml) induced THP1 macrophages (Figure 4A–4C). Furthermore, Lipidom treatment also significantly (p <0.001) reduced the NLRP3 inflammasome involved in the progression of atherosclerosis [7] as observed by the decrease in levels of LPS and ATP induced IL-1β cytokine release (Figure 4D).

**Figure 4.**
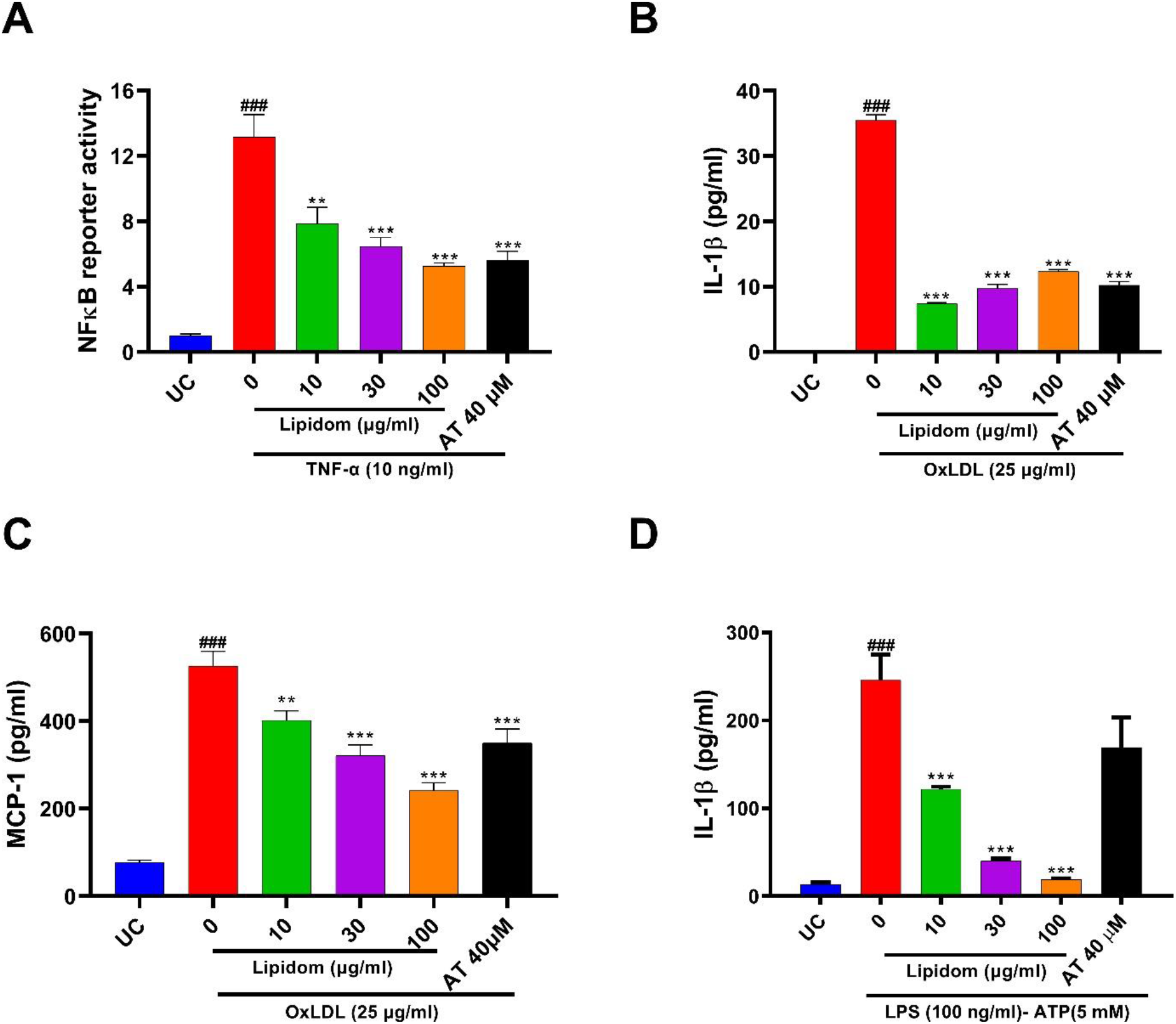
Lipidom possesses anti-inflammatory activity. **A)** Lipidom decreased the TNF-α (10 ng/ml) induced (24 hr) THP1-Blue NF-κB reporter cells. **B)** Lipidom decreased the IL-1β cytokine release from the OxLDL (25 μg/ml) induced differentiated (PMA 60 ng/ml) THP1 macrophages (72 hr). **C)** Lipidom decreased the MCP-1 chemokine release from the OxLDL (25 μg/ml) induced differentiated THP1 macrophages (72 hr). **D)** Lipidom decreased the NLRP3 inflammasome mediated IL-1β cytokine release from differentiated (PMA 20 ng/ml) THP1 macrophages.

### Lipidom decreased the ROS generation and Lipid accumulation in OxLDL induced THP1 macrophages

Treatment with Lipidom (100 μg/ml) decreased the ROS generation and Lipid accumulation in OxLDL (25 μg/ml) induced THP1 cells as observed by the fluorescence microscopy images (Figure 5A). Furthermore the % ROS levels were quantified wherein we observed a dose dependent decrease (Figure 5B). The lipid peroxidation levels as represented by the amount of MDA in cells were also decreased by Lipidom treatment (Figure 5C).

**Figure 5.**
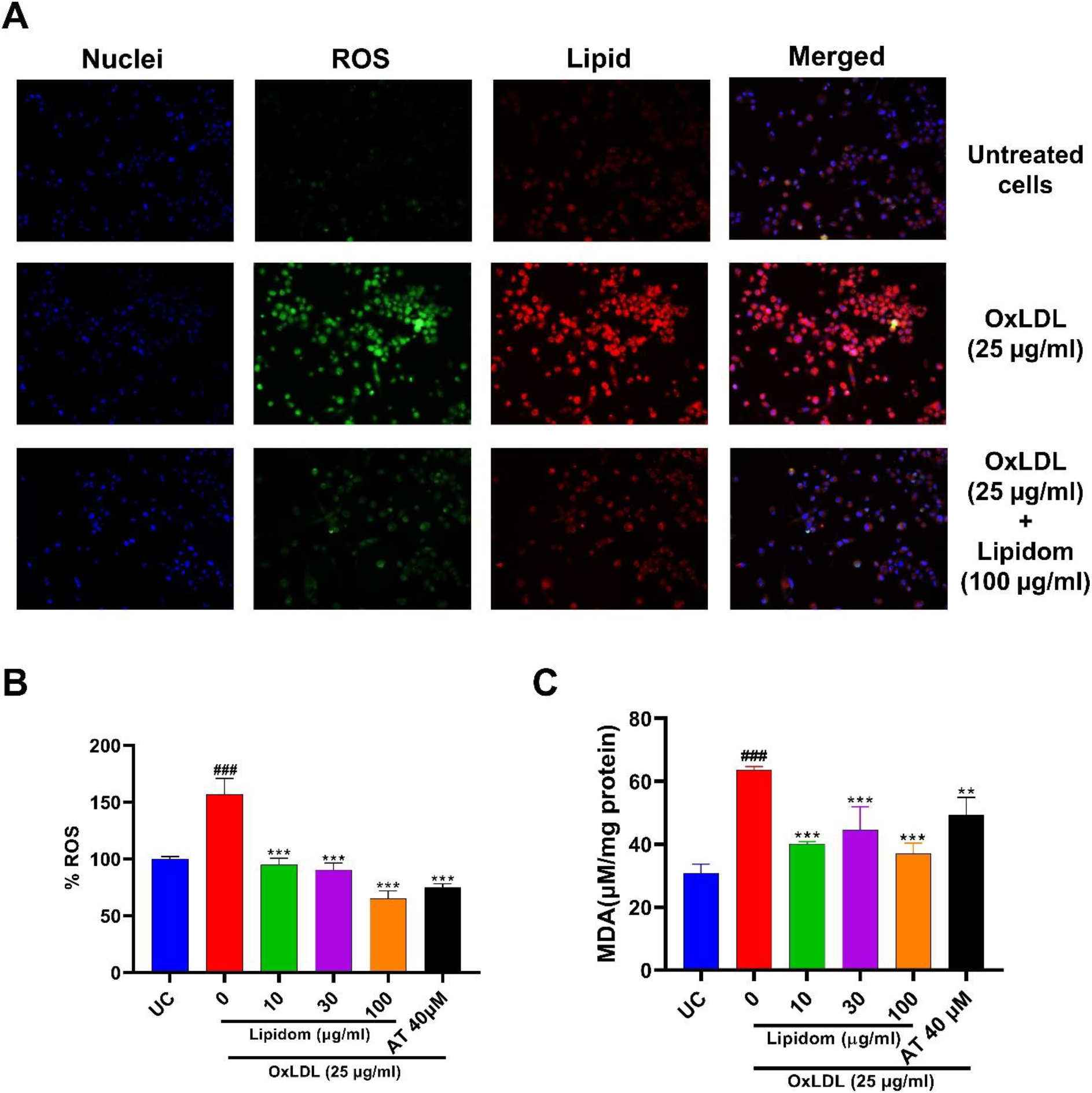
Lipidom decreased Lipid accumulation, ROS and MDA levels. **A)** Lipidom decreased the lipid (red) and ROS (green) accumulation in OxLDL (25 μg/ml) induced differentiated THP1 macrophages (72 hr). Magnification= 200×. **B, C)** Lipidom decreased the ROS and MDA levels in OxLDL (25 μg/ml) induced differentiated THP1 macrophages (72 hr).

### Lipidom the levels of genes involved in the foam cell formation

Lipidom is involved in the modulation of genes responsible for the progression of atherosclerosis and development of chronic inflammatory conditions. Genes namely TNF-α, MCP-1 (inflammation), FAS, ABCG1 (lipid homeostasis), Nrf-2 (oxidative stress), and FGF21 (lipid metabolism) are modulated by Lipidom in a dose-dependent manner (Figure 6A–6F).

**Figure 6.**
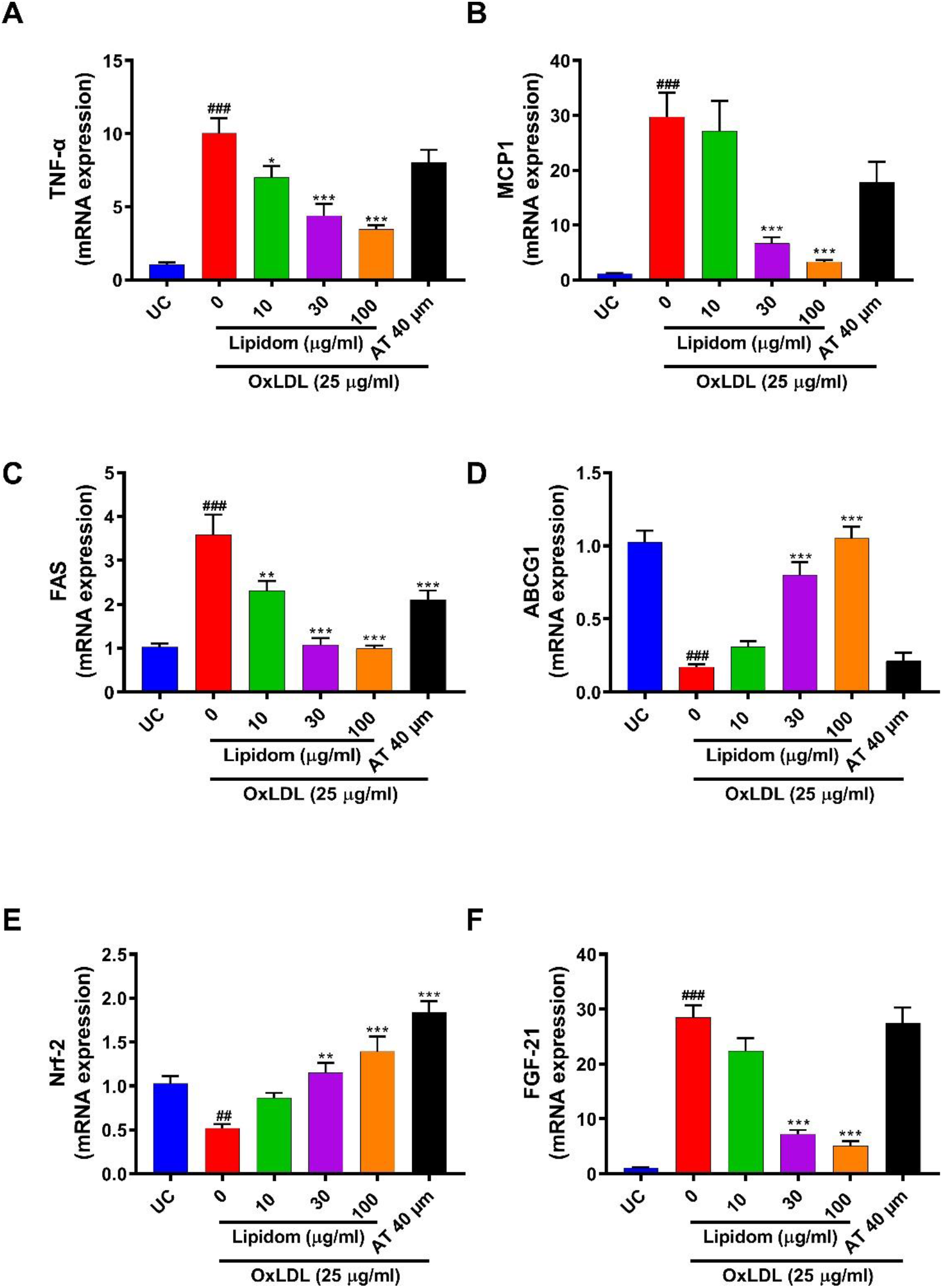
Lipidom modulated mRNA expression in OxLDL induced THP1 macrophages. Lipidom modulated the levels of **A)** TNF-α, **B)** MCP-1), **C)** FAS, **D)** ABCG1, **E)** Nrf-2, and **F)** FGF-21 genes involved in atherosclerosis development and progression.

### Lipidom treatment decreased lipid and triglyceride levels *C. elegans*

Treatment with Lipidom (3, 10, and 30 μg/ml) significantly (p <0.001) reduced the lipid and triglyceride accumulation (Figure 7C, 7D) in *C. elegans*, without affecting nematode reproduction (brood size) and growth (% Adult) (Figure 7A, 7B). These results suggest that Lipidom is able to modulate the lipid metabolism without hindering the normal growth and reproduction cycle of *C. elegans*.

**Figure 7.**
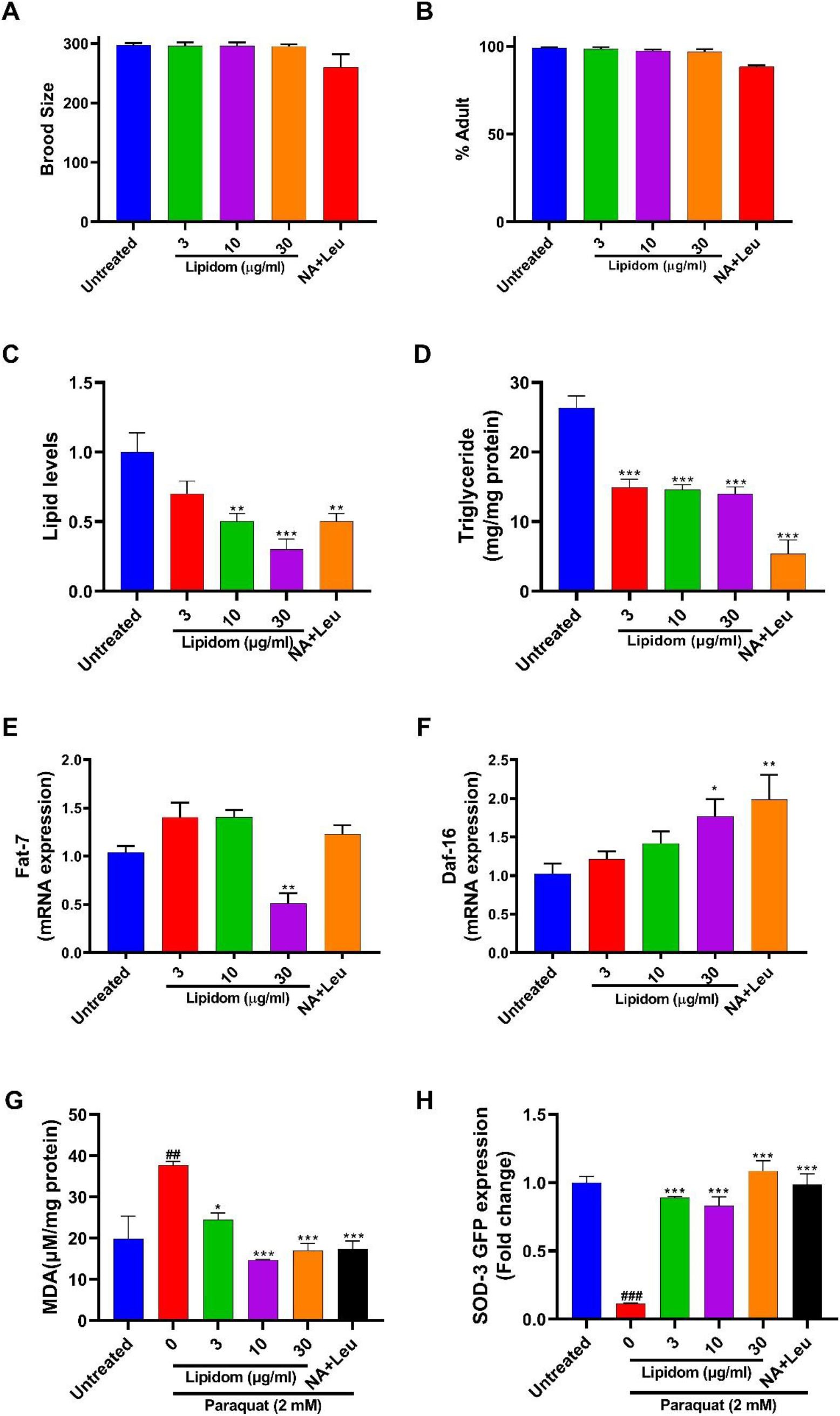
Lipidom showed lipid lowering and anti-oxidant activity on *C. elegans*. Lipidom showed no adverse effect on **A)** brood size and **B)** % Adult survival. Lipidom decreased **C)** lipid and **D)** triglyceride levels in *C. elegans*. Lipidom modulated **E)** Fat-7 and **F)** Daf-16 genes involved in fat metabolism. **G)** Lipidom decreased the MDA levels in N2 strain of *C. elegans*. **H)** Lipidom increased the SOD-3 GFP levels in CF1553 (muls84[pAD76(sod-3::GFP)]) strain of *C. elegans*.

### Lipidom treatment modulated the Fat-7 and Daf-16 gene expression in *C. elegans*

Lipidom treatment significantly (p <0.01) downregulated the stearoyl-CoA desaturase gene (Fat-7) (Figure 7E) thereby further aiding the decrease of lipid accumulation inside the nematode. Moreover, Lipidom treatment also significantly (p <0.05) enhanced the expression of Daf-16 (Figure 7F), the sole *C. elegans* forkhead box O (FOXO) homologue which is known to regulate lipid metabolism by targeting fatty acid desaturation [8]. Taken together, it was observed that Lipidom aids in the decrease of lipid accumulation as observed by its effect on the expression levels of genes involved in lipid metabolism.

### Lipidom treated modulated the oxidative stress levels of paraquat induced *C. elegans*

Lipidom (3, 10, and 30 μg/ml) treatment significantly (p <0.001) decreased the MDA levels (Figure 7G) of the worms stimulated with Paraquat (2 mM). Furthermore, SOD-3 expression of the CF1553 strain of *C. elegans* was significantly (p < 0.001) increased post Lipidom treatment as observed from the increase in the levels of their fluorescent GFP levels. As, SOD-3 is involved in encoding of mitochondrial SOD, a well-known free radical scavenging SOD; the increase in SOD-3 expression by Lipidom signifies that it has potent anti-oxidant properties.

## Discussion

Cardiovascular diseases account for 17.9 million deaths every year, majority of which are due to atherosclerosis. Chronic inflammation is a major risk factor of atherosclerosis progression. Currently. the pharmacotherapy relies on the control of inflammation by use of non-steroidal anti-inflammatory drugs (NSAIDs), coxibs and HMG CoA reductase inhibitors (statins) but due to increased risk of life-threatening adverse effects like muscle pain, myocardial infarction, stroke, and rhabdomyolysis their use becomes limited [9, 10]. Over the past few decades, herbal medicines have gained traction as feasible therapeutic agents for prevention and treatment of atherosclerosis due to their potential of targeting multiple steps involved in pathogenesis and lack of adverse effects [11].

The current study aimed to analyse the phytochemistry and bioactivity of the Ayurvedic prescription medicine Lipidom. The phytometabolite characterization revealed the presence of Gallic acid, Protocatechuic acid, Corilagin, Ellagic acid, Cinnamic acid, Guggulsterone E, Guggulsterone Z in Lipidom. These phytometabolites are known to possess anti-oxidant, anti-inflammatory and lipid lowering properties [12–17]. In order to evaluate the pharmacological activities of these phytochemicals we determined the effect of Lipidom on OxLDL induced THP1 macrophage in-vitro. Furthermore, we also evaluated its therapeutic potential via *C. elegans* based in-vivo model.

Prior to the evaluation of the bioactivity of Lipidom we evaluated its cytosafety on THP1 macrophages. Lipidom was found to be cytosafe at all physiologically relevant concentrations. Furthermore, the relevant concentration of OxLDL for inducing foam cell like properties in THP1 cells were also determined. The concentration which showed least toxic effect was selected. The cytosafety evaluations allowed us to rule out any confounding bias in our obtained results. As, inflammation is the primary reason for atherosclerosis development and progression we evaluated Lipidom against various mediators of inflammation. The TNF-α driven activation of NFκB is involved in regulation of inflammatory cascade at various stages of atherosclerosis, right from plaque formation to its stabilization and rupture [18, 19]. Lipidom treatment was found to decrease the TNF-α induced NFκB activity in a concentration dependent manner.

Besides NFκB another inflammatory mediator IL-1β is also involved in multiple stages of atherosclerosis. IL-1β induces an inflammatory response in endothelial cells and enhances expressions of adhesion factors and chemokines, and promotes the accumulation of inflammatory cells in blood vessels and their invasion into the local intima of blood vessels, which often happens at the initiation of atherosclerosis. One of the stimulated chemokine is monocyte chemoattractant protein (MCP-1) which recruits mononuclear phagocytes and is closely related to atherosclerosis [20]. It was observed that Lipidom treatment significantly lowered the release of both IL-1β and MCP-1 from the OxLDL induced THP1 macrophages. A key feature of atherosclerosis is development of inflammation with macrophage infiltration which is driven by NLRP3 inflammasomes. The NLRP3 inflammasomes regulate caspase-1 activation and subsequent processing of pro-IL-1β, trigger vascular wall inflammatory responses and lead to progression of atherosclerosis [21]. Lipidom treatment also decreased NLRP3 driven IL-1β cytokine release from THP1 macrophages upon dual triggering by LPS and ATP. Hence, Lipidom possess a wide array of anti-inflammatory properties which helps in blocking the progression of atherosclerosis.

The accumulation lipid inside macrophages leads to foam cell formation, a hallmark of atherosclerotic lesions. The formation and retention of lipid loaded macrophages in these lesions exacerbates the progression of atherosclerosis [22]. The formation of atherosclerotic plaque also increases due to oxidative stress [23]. Lipidom was able to decrease lipid accumulation in THP1 macrophages as observed by fluorescence microscopy of OxLDL induced cells. Furthermore, Lipidom treatment decreased ROS and MDA formation. Taken together it was observed that Lipidom possesses potent anti-oxidant and lipid lowering activities.

In order to decipher the mode of action of Lipidom we evaluated the gene expression of OxLDL induced THP1 macrophages and found that Lipidom acts on the pathways responsible for inflammation (TNF-α, MCP-1), lipid accumulation (FAS, ABCG1, FGF-21) and oxidative stress (Nrf2) [24–26]. It was found that Lipidom was able to normalize the expression of these genes by a multifaceted mode of action.

The efficacy of Lipidom treatment was also observed in the in-vivo nematode model. Lipidom treated *C. elegans* did not show any defects in survival and reproduction. Moreover, Lipidom decreased the lipid accumulation and triglyceride levels of the nematodes by regulation of Fat-7 and Daf-16 genes which are involved in lipid metabolism. The antioxidant potential of Lipidom was also evaluated on the nematode where it was observed that in presence of an oxidative stress inducing agent like paraquat the worms treated with Lipidom were able to normalize MDA formation. Also, Lipidom treatment increased the SOD-3 expression in the mutant *C. elegans* strain CF1553. As, SOD-3 is involved in defence against oxidative stress during atherosclerosis [27, 28] this potential of Lipidom can play an important role in preventing the progression of plaque formation.

In conclusion, Lipidom was able to show experimental evidences against lipid accumulation disorders, as observed from its anti-inflammatory, lipid lowering and anti-oxidant properties, by targeting multiple pathways responsible for initiation and progression of disease aetiology. Taken together, our study suggests that Lipidom has potential pharmacological properties against cardiovascular disorders like dyslipidaemia and atherosclerosis.

## Author Contributions

AB: Conceptualization, Planning, Visualization, Supervision. VG: Conceptualization, Planning, Visualization, Methodology, Investigation, Data curation, Formal analysis, Writing original draft. NP: Methodology, Investigation, Formal analysis. RS: Methodology, Investigation, Formal analysis. MT: Methodology, Investigation, Formal analysis. MR: Methodology, Investigation, Formal analysis. RD: Data curation, Writing - review & editing, Visualization, Project administration, Supervision. AV: Writing - review & editing, Project administration, Conceptualization, Visualization, Supervision.

## Acknowledgments

We extend our gratitude to Ms. Deepika Rajput for her support in the biochemical analysis. We are also grateful to Mr. Tarun Rajput, Mr. Gagan Kumar, and Mr. Lalit Mohan for their swift administrative support.

